# An open-hardware platform for optogenetics and photobiology

**DOI:** 10.1101/055053

**Authors:** Karl P. Gerhardt, Evan J. Olson, Sebastian M. Castillo-Hair, Lucas A. Hartsough, Brian P. Landry, Felix Ekness, Rayka Yokoo, Eric J. Gomez, Prabha Ramakrishnan, Junghae Suh, David F. Savage, Jeffrey J. Tabor

## Abstract

In optogenetics, researchers use light and genetically encoded photoreceptors to control biological processes with unmatched precision. However, outside of neuroscience, the impact of optogenetics has been limited by a lack of user-friendly, flexible, accessible hardware. Here, we engineer the Light Plate Apparatus (LPA), a device that can deliver two independent 310 to 1550 nm light signals to each well of a 24-well plate with intensity control over three orders of magnitude and millisecond resolution. Signals are programmed using an intuitive web tool named Iris. All components can be purchased for under $400 and the device can be assembled and calibrated by a non-expert in one day. We use the LPA to precisely control gene expression from blue, green, and red light responsive optogenetic tools in bacteria, yeast, and mammalian cells and simplify the entrainment of cyanobacterial circadian rhythm. The LPA dramatically reduces the entry barrier to optogenetics and photobiology experiments.

---

In 2005, Deisseroth and co-workers reported the expression of a blue light activated microbial ion channel (opsin) in mammalian neurons and its use for physiologically-relevant, millisecond timescale optical control of neural activity *in vitro*^1^. However, because no instrument existed for delivering the necessary intensity of light to specific brain regions in live animals without major side effects, optogenetics contributed few neurobiological insights between 2005 and 2009^2^. During this period, the group engineered the optical neural interface (ONI) — a brain-implantable optical fiber with a laser diode light source. By combining the ONI with microbial opsins, they achieved optogenetic control of rat motor behavior^3^ and identified neural activity patterns that cause sleeping mice to wake^4^. The ONI was rapidly adopted by the greater community and combined with opsins and other photoreceptors, resulting in a wave of breakthroughs in a short time period^2^.

Following Quail and co-workers’ pioneering engineering of a red/far red light-switchable *S. cerevisiae* transcriptional regulatory (promoter) system in 2002^5^, photoreceptors with diverse spectral properties have been used to control a remarkable range of cell biological processes in mechanistically tractable model organisms as well. For example, light-switchable promoter systems have been engineered in *E. coli*^6,11^, cyanobacteria^12^, yeast^13,14^, mammalian cells^15–, 24^, fruit flies^25^, zebrafish^21, 26^ and plants^27^. Translation^28, 29^, proteolysis^30^, membrane recruitment^13, 31^, signaling^31–39^, ER-to-cytoplasm^40^ and nuclear^41–44^ translocation, and genome editing^13,45,46^ have also been placed under optogenetic control.

However, no optical hardware has been developed to enable the broad research community to properly utilize these non-neural optogenetic tools, limiting their impact. For example, we recently engineered the Light Tube Array (LTA), a light emitting diode (LED)- based device that exposes 64 shaking incubated culture tubes to programmable light signals with an intensity range over three orders of magnitude and millisecond resolution^47^. Though the LTA enables unrivaled control of gene expression dynamics^47^, construction requires custom machined components, specialized assembly tools, and knowledge of electronic system design and programming is done in computer language. Additionally, experiments are not scalable due to the large instrument size (0.02 m^3^) and requirement for connection to an external computer. Furthermore, the tubes are costly (~$0.10/each) and restrict experiments to suspension culture organisms such as bacteria and yeast. Finally, the LEDs are permanently soldered, limiting optical flexibility. In another example, Moglich and coworkers modified the injection port of a Tecan microplate reader with an optical fiber and eight LEDs from 385 – 850 nm^48^. While this clever design enables programmable sample illumination via the commercial software, the Tecan instrument costs ~$40,000. Other recent designs^35,42,49–52^ suffer various limitations and have not found widespread adoption.

We have designed the LPA for compatibility with a wide range of optogenetics and photobiology experiments and model organisms, low device and consumable costs, high scalability and throughput, and accessibility by laboratories without hardware expertise. We demonstrate its capabilities by recapitulating and extending our LTA results in *E. coli*, performing yeast and mammalian optogenetic gene expression control experiments, and easing the entrainment of cyanobacterial circadian rhythm. Additionally, because the LPA and Iris are open source, they may be freely modified and extended by the community for additional functionalities.

## Results

### LPA design and assembly

The core of the LPA is a printed circuit board (PCB) (**Supplementary Fig. 1a-c**) outfitted with a Secure Digital (SD) card reader (**Supplementary Figs. 1c** and **2**), an Atmel ATMega328a microcontroller (**Supplementary Figs. 1c** and **3**), 3 LED drivers (**Supplementary Figs. 1c** and **4**), 48 solder-free LED sockets (**Supplementary Fig. 1a**), a power regulating circuit (**Supplementary Fig. 5**), and other standard electronics components (**Supplementary Table 1**). The only external connection is to a 5v power supply (**Fig. 1a** and **Supplementary Fig. 6**). The unpopulated PCB can be ordered from a commercial supplier using the provided fabrication files (**Supplementary Information**). The PCB can be assembled (populated with electronic components) via do-it-yourself (DIY) (**Supplementary Procedure 1** and **Supplementary Table 2**) or commercial (**Supplementary Procedure 1**) soldering procedures. We recommend DIY installation of the LED sockets using a 3D printed socket alignment tool (**Supplementary Fig. 7a** and **Supplementary Procedure 1**) to ensure consistent geometries and illumination across the wells. We have provided a video tutorial demonstrating PCB assembly (**Supplementary Video 1**).

**Figure 1.**
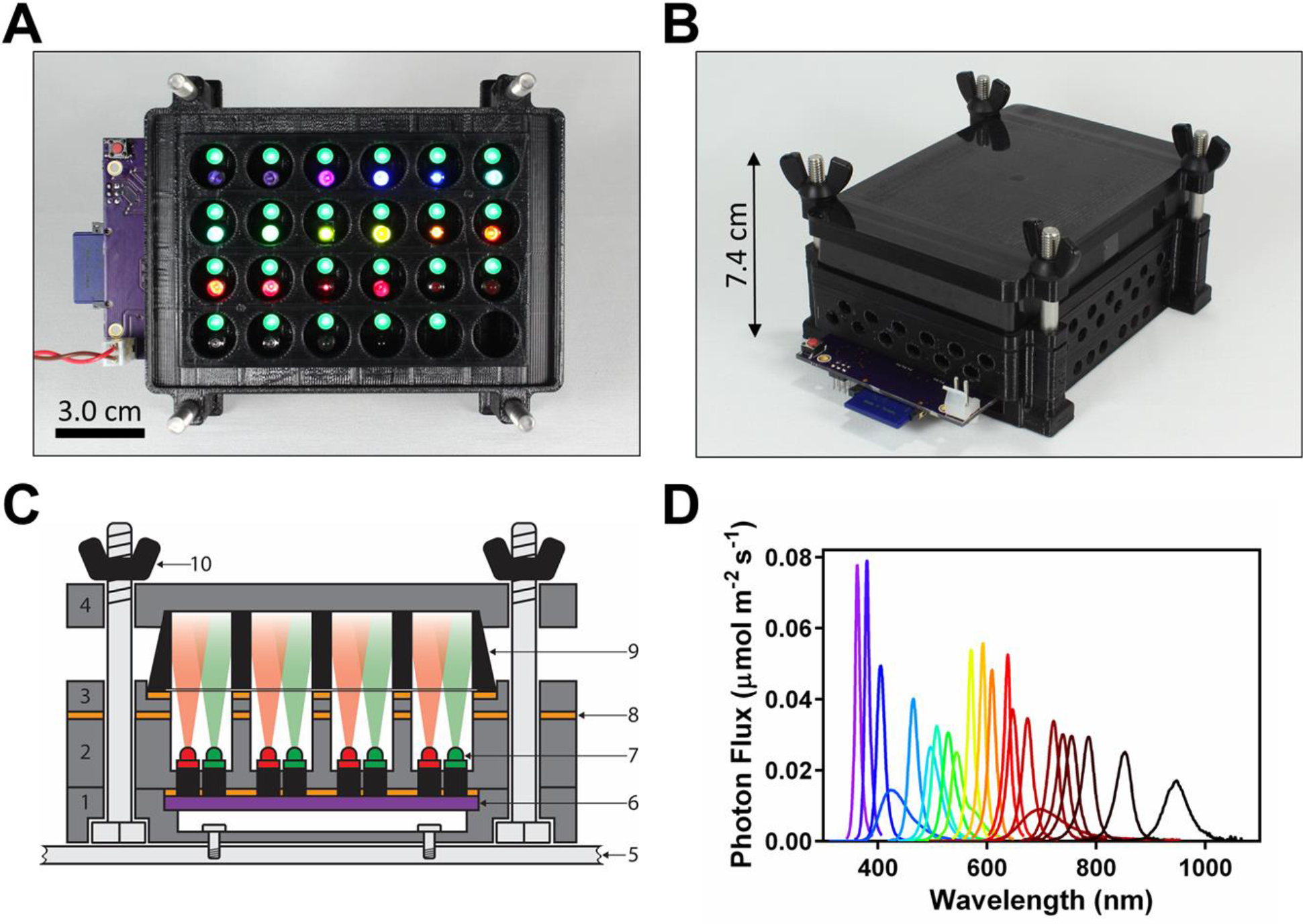
The LPA. (**a**) Top-down view of an LPA lacking the 24-well plate and plate lid. In the configuration shown, the device contains a 535 nm LED in each top position and a variable wavelength LED (364–947 nm) in each bottom position. The SD card (blue), circuit board (purple), reset button (red) and power cord (red/black wires) are visible. (**b**) Fully assembled LPA. (**c**) Schematic cross-section showing the light path from the LEDs to the culture plate wells. (1) mounting plate, (2) LED spacer, (3) plate adapter, (4) plate lid, (5) PCB, (6) LED atop LED socket, (7) gasket, (8) 24 well plate, (9) wing nut assembled with mounting bolt. (d) Spectra of 22 LEDs (**Supplementary Table 5**) used in this study measured and calibrated using a spectrophotometer and probe adapter (**Supplementary Figs. 6b, 20**, and **Supplementary Procedure 8**).

The microcontroller converts an Iris-generated file stored on the SD card to LED driver control signals. Each LED driver individually regulates the intensity of 16 LEDs over 4096 levels with a 1 ms refresh rate via 12-bit pulse width modulation of output current up to 20 mA. The output of each driver, and thus LED intensity, can be further controlled with 6-bit resolution by adjusting dot correction settings in the device’s firmware (**Supplementary Figs. 8–14, Supplementary Tables 3 and 4, Supplementary Procedures 2** and **3**, and **Supplementary Note 1**).

Each socket secures a 5 mm through-hole LED via a friction fit (**Supplementary Fig. 15**). These LEDs are commercially available with wavelengths from 310 to 1550 nm (**Supplementary Table 5**), enabling plug-and-play optical reconfiguration (**Supplementary Procedure 4**). The 20 mA driver output permits full range control of most 5 mm LEDs.

A 3D printed chassis houses the assembled PCB and 24-well plate (**Fig. 1a–Fig. 1c**). The chassis comprises i) a mounting plate, ii) LED spacer, iii) plate adapter, and iv) plate lid (**Supplementary Fig. 16a-d**). We have designed 4 mounting plates, ensuring compatibility with most shaker platforms (**Supplementary Information**). The mounting plates contain recessed holes (**Fig. 1c** and **Supplementary Fig. 16a**) for upward-facing bolts that align and stack the remaining 3 modules. The LED spacer (**Supplementary Fig. 16b**) creates a fixed distance between each LED socket pair and overhead well, and optical isolation between wells. The spacer is mated with the PCB and the resulting assembly stacked atop the mounting plate. The plate adapter (**Supplementary Fig. 16c**) is placed atop the spacer. A black-walled, transparent plastic bottomed 24 well plate (**Supplementary Fig. 17** and **Supplementary Table 6**) is aligned and held in place by the plate adapter. An adhesive foil plate cover (**Supplementary Table 6**) provides a sealed environment for each well. Laser cut nitrile gaskets (**Supplementary Fig. 18** and **Supplementary Procedure 5**) are placed at each of the 3 interfaces above the PCB to reduce optical contamination. Finally, the lid (**Supplementary Fig. 16d**) is placed atop the plate and the complete assembly is secured with wing nuts (**Fig. 1b** and **Supplementary Table 6**). All chassis modules can be 3D printed using supplied files (**Supplementary Information, Supporting Procedure 6**). We have provided a written procedure and instructional video demonstrating LPA assembly (**Supplementary Procedure 7** and **Supplementary Video 2**).

### LPA programming with Iris

Iris can be freely accessed at http://iris.taborlab.rice.edu or run locally using the provided source code (**Supplementary Note 2**). Light programs are specified in Steady State, Dynamic, or Advanced Modes (SSM, DM, AM). In SSM, time-invariant intensities for each LED are entered within an Input Panel (**Fig. 2a, left**). In DM, constant, step, sinusoidal, or piecewise light signals are specified using parameters such as initial and final intensity, time of step, or wave period, amplitude, and offset (**Supplementary Note 3**). The same signal is run by all 24 top LEDs, and the same, or a second signal can be run by all 24 bottom LEDs. AM (**Fig. 2b, left**) implements DM signals via our recent staggered-start protocol, which allows kinetic experiments to be performed with far less sample handling^47^ (**Supplementary Fig. 19, Supplementary Table 7**). The light signal run in each well can be randomized (**Supplementary Note 4**) to mitigate any well-to-well variability. We have provided videos demonstrating Iris light signal programming in all 3 modes (**Supplementary Videos 3–5**).

**Figure 2.**
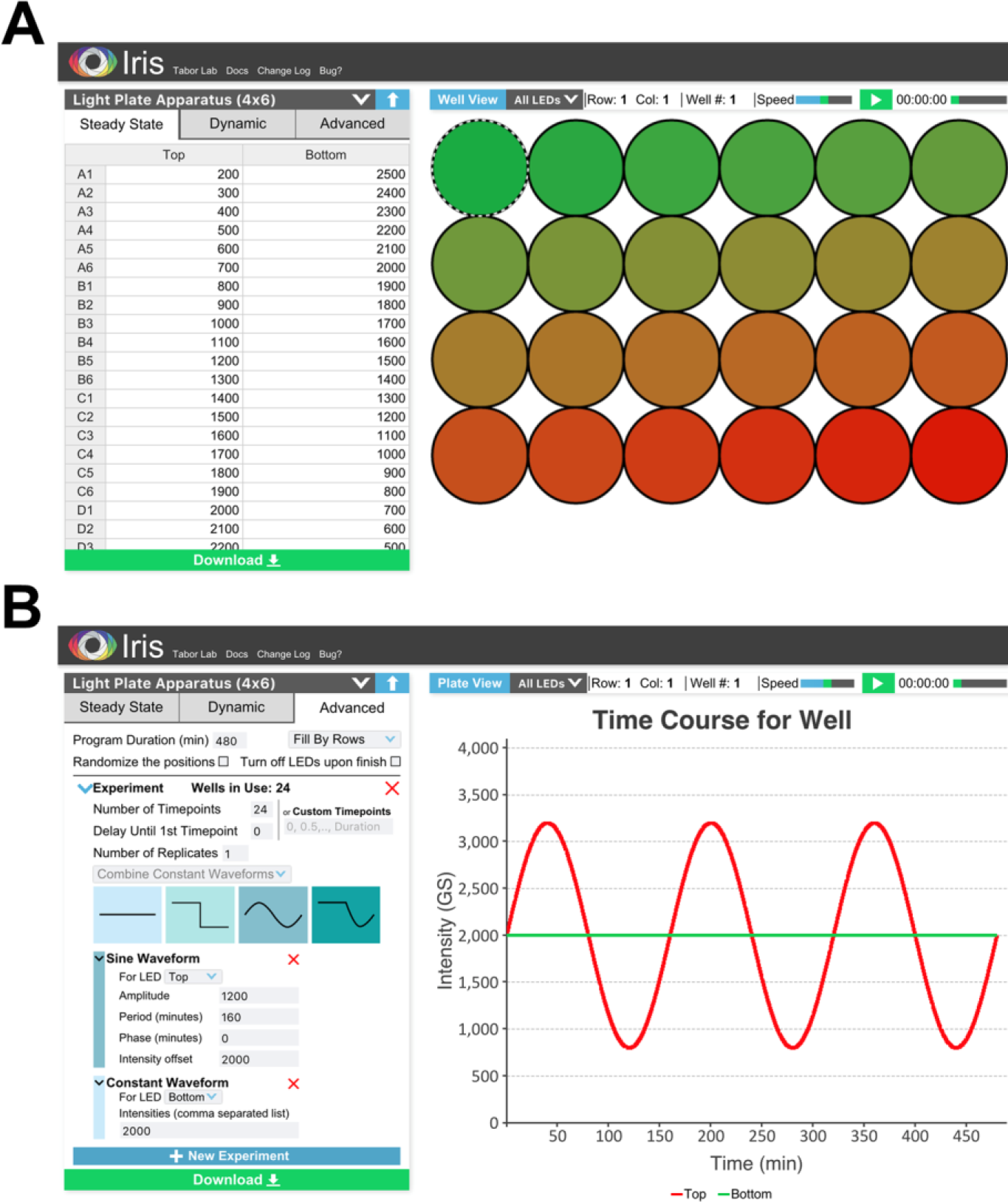
Iris. (**a**) Steady State Mode. Constant light functions are specified by entering the desired greyscale intensity (0–4095) of up to all 48 LEDs in the Input Panel spreadsheet (left). The download button is shown at the bottom of the Input Panel. In Plate View mode (shown), the Simulation Panel (right) displays a schematic visualization of the output intensity of all 48 LEDs. The top and bottom LEDs are visualized as red and green, respectively, regardless of the actual LEDs used. (**b**) Advanced Mode. Constant and dynamic light functions are specified in the Input Panel. In the latter case, Iris automatically runs a staggered-start algorithm (**Supplementary Fig. 19**). In Well View mode (shown), the Simulation Panel (right) visualizes the output of the top and bottom LEDs in a given well over the duration of the experiment. The Simulation Panel also plays movies of specified light function

Light programs can be viewed and debugged at the plate (**Fig. 2a, right**) or individual well (**Fig. 2b, right**) levels using the Simulation Panel. When the program is satisfactory, the download button (**Fig. 2a, b**) is pressed. Iris then generates i) a device-readable binary file (.lpf, **Supplementary Table 8**) used to run the LPA, ii) a session file for reloading a program into Iris at a later time, and iii) a CSV file containing user-readable well randomization information in a zip folder. The .lpf is then transferred to an SD card, which is inserted into the LPA, and the Reset Button (**Supplementary Fig. 1a**) is pressed to run the light program. Included Python scripts can be used to automate .lpf design (**Supplementary Information, Supplementary Note 5**), increasing throughput.

### LPA calibration

We have developed a simple method utilizing a gel imager and MATLAB script to calibrate the outputs of multiple LPA LEDs of one spectrum to one another (**Supplementary Information, Supplementary Procedure 8**). To calibrate absolute outputs, or the outputs of LEDs with different spectra, we have developed a method combining a probe spectrophotometer and 3D printed adapter that fits the LPA wells (**Supplementary Figs. 6b, 20 and Supplementary Procedure 8**). Using either method, outputs are adjusted by loading two manually generated text files specifying the grayscale and dot correction values for each LED (**Supplementary Procedure 8**, examples in **Supplementary Information**) on to the SD card before running an experiment. Calibration typically reduces LEDs output variability to <1% (**Supplementary Fig. 21**).

### Benchmarking the LPA with *E. coli* CcaS-CcaR

CcaS-CcaR is a green light activated, red de-activated two component system (TCS) that we previously engineered to control gene expression in *E. coli* (**Fig. 3a**)^8^. Using the LTA and a superfolder GFP (sfGFP) output, we characterized the relationship between green and red light inputs and transcriptional output dynamics (i.e. I/O)^47^. We developed a predictive mathematical model of the I/O, which we used to pre-compute green light time courses to drive CcaS-CcaR output to follow tailor-made (reference) gene expression signals with high predictability^47^.

**Figure 3.**
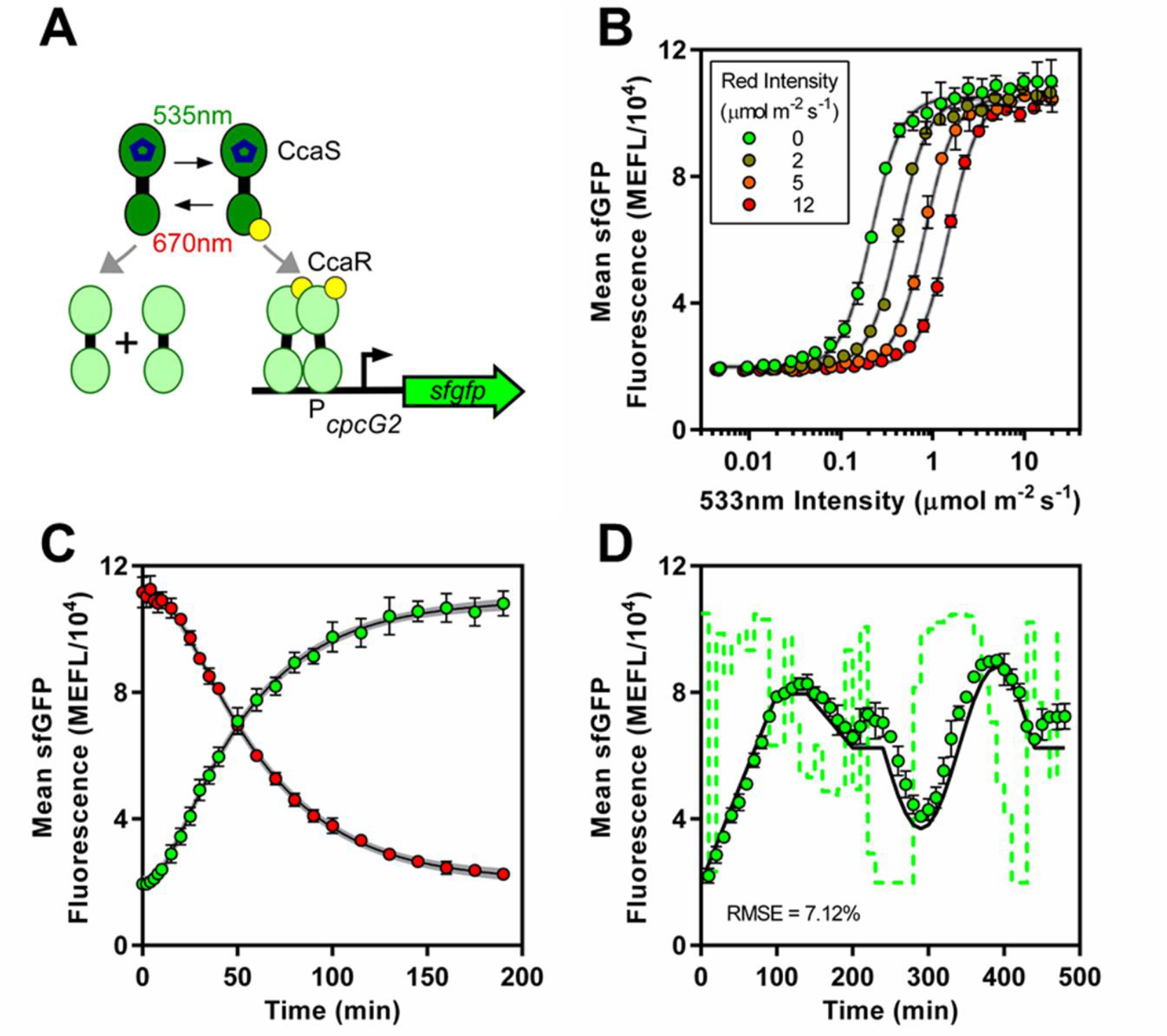
Benchmarking the LPA against *E. coli* CcaS-CcaR. (**a**) CcaS-CcaR system. (**b**) CcaS-CcaR 533 nm light intensity versus sfGFP transfer functions in the presence of increasing 678 nm. (**c**) Kinetic response to an increase in 533 nm light from 0.00 to 17.88 μmol m^−2^ s^−1^ and simultaneous decrease in 678 nm light from 12.79 μmol m^−2^ s^−1^ to 0.00 μmol m^−2^ s^−1^ (green dots), or decrease in 533 nm light from 17.88 to 0.00 μmol m^−2^ s^−1^ and simultaneous increase in 678 nm from 0.00 μmol m^−2^ s^−1^ to 12.79 μmol m^−2^ s^−1^ (red dots). Black lines represent best fits (**Supplementary Table 9** and **10**) of our previous CcaS-CcaR mathematical model^47^ to these data. Gray envelopes represent 95% confidence intervals. (**d**) Biological Function Generation. Reference waveform (black line), precomputed 533 nm light intensity time course (green dashed lines), experimental sfGFP levels (green dots). Constant 12.79 μmol m^−2^ s^−1^ 678 nm was applied. 533 nm intensity values are mapped to sfGFP units through the transfer function in panel a. RMSE between experimental data and reference over three days is shown. Error bars represent the SEM of three experiments over three days.

We aimed to benchmark the LPA by repeating these CcaS-CcaR experiments. First, we measured the steady state response to green light intensity by outfitting the bottom positions of an LPA with 533 nm (green) LEDs and programming them to emit between 0.00 and 20.10 μmol m^−2^ s^−1^ photons. We grew our previous strain in a shaking LPA under these conditions and measured sfGFP output using flow cytometry (**Methods**). We observed that sfGFP increases sigmoidally with green intensity with a 5.3-fold dynamic range and Hill parameter (*n_H_*) = 2.6 (**Fig. 3b**), tightly consistent with LTA measurements. The intensity resulting in half maximal output (*k_1/2_*) is 0.22 μmol m^−2^ s^−1^, 4.8-fold greater than the LTA value. This discrepancy is likely due to the shorter length between LED and sample and small probe size. Because LED emission is conical (**Fig. 1c**), this length difference concentrates photons on the probe, artificially elevating measured intensity values. This discrepancy could be eliminated using improved probe designs.

Next, we benchmarked the effect of red light by outfitting the top and bottom positions with 533 and 678 nm (red) LEDs and re-measured the green light response in the presence of 2.00 – 12.00 μmol m^−2^ s^−1^ red photons. We observed that dynamic range and *n_H_* remain unchanged while *k_1/2_* increases by 0.11 μmol m^−2^ s^−1^ per 1.00 μmol m^−2^ s^−1^ red light (**Fig. 3b** and **Supplementary Table 9**), consistent with our previous data.

Finally, we used the LPA to characterize the kinetic response of CcaS-CcaR to step changes in green intensity. We observed a 4.5 min delay, an 11 min transcription rate switching time, and sfGFP switching dynamics set by the cell division time (**Fig. 3c**), equivalent to LTA results. We used these data to re-calibrate the parameters of our model to the LPA conditions (**Methods** and **Supplementary Table 10**). Finally, we successfully used the model to program sfGFP expression to follow a challenging waveform comprising linear ramps, a hold, and a sine wave with high predictability (**Fig. 3d**). We conclude that the LPA meets the same performance standards at the LTA.

### Characterizing CcaS-CcaR forward and reverse action spectra

The action spectrum is the relationship between input wavelength and output activity. While all optogenetic tools have forward (ground state) action spectra (FAS), photoreversible tools also have reverse (activated state) action spectra (RAS).

We next aimed to demonstrate that the LPA can be used to characterize the FAS and RAS of optogenetic tools using CcaS-CcaR as a model. To this end, we outfitted the bottom position of a device with LEDs of output wavelength between 364 and 947 nm (**Fig. 1d** and **Supplementary Table 5**) and programmed each to emit 0.40 μmol m^−2^ s^−1^, or 2\**K_1/2_* for the 533 nm LED (**Fig. 3b**). We then measured the steady state response of CcaS-CcaR to each input as before. This experiment reveals a maximal activating wavelength of 533 nm and >50% maximal activation between 533 and 571 nm (**Fig. 4**). We previously generated a course-grained map of the CcaS-CcaR FAS using six wavelengths with a rudimentary instrument in an experiment that required approximately one week^8^. By contrast, the LPA enables us to measure a 24 wavelength FAS in a single day.

**Figure 4.**
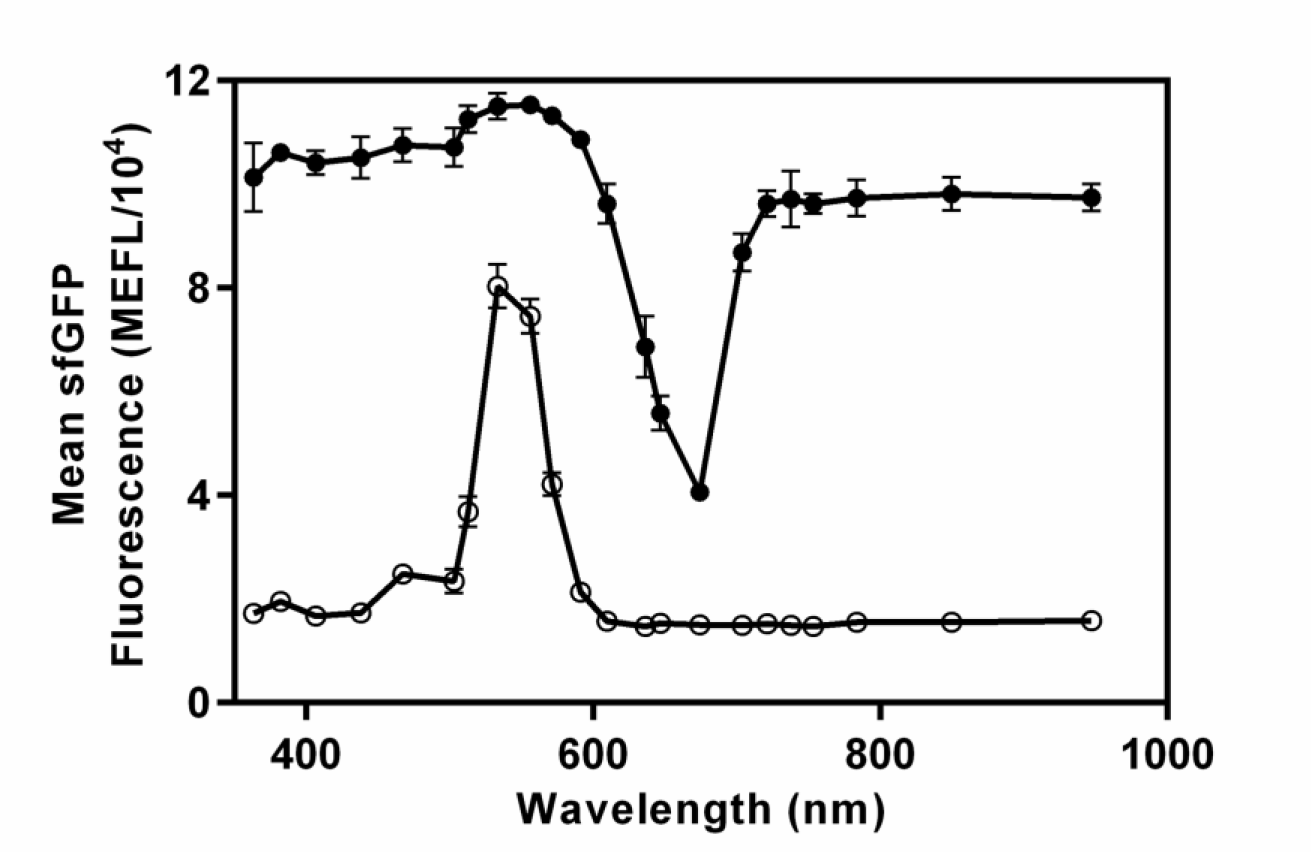
Using the LPA to characterize CcaS-CcaR forward and reverse action spectra. For the FAS (hollow circles), bacteria were exposed to 0.397 μmol m^−2^ s^−1^ photons from a variable wavelength bottom LED (**Fig. 1a,c**). For the RAS (black circles), bacteria were exposed to 0.397 μmol m^−2^ s^−1^ from the maximally activating 533 nm LED in the top position and 3.21 μmol m^−2^ s^−1^ photons from the variable wavelength bottom LED. Error bars represent the SEM of three experiments over a single day.

To characterize the RAS, we outfitted the top positions of a device with the 533 nm LED and the bottom positions with the same LEDs used in the FAS experiment. We set the 533 nm LED to 0.40 μmol m^−2^ s^−1^ and each bottom LED to 3.21 μmol m^−2^ s^−1^ because CcaS-CcaR has approximately 10-fold greater sensitivity to green than red light (**Fig. 3b**). Consistent with *in vitro* absorbance measurements of activated CcaS^53^, exposure of our CcaS-CcaR expressing strain to these inputs reveals maximal deactivation by the 678 nm LED (**Fig. 4**). To our knowledge, this is the first measurement of the RAS of an optogenetic tool.

### Characterizing the *S. cerevisiae* CRY2-CIB1 yeast two hybrid system

Tucker and coworkers previously engineered a blue light activated *S. cerevisiae* promoter system by fusing Cryptochrome 2 (CRY2) to the Gal4 DNA binding domain (DBD) and the interaction domain of its blue-light dependent heterodimerizing partner CIB1 to the Gal4 activation domain in a yeast two-hybrid (Y2H) system^13^ (**Fig. 5a**).

**Figure 5.**
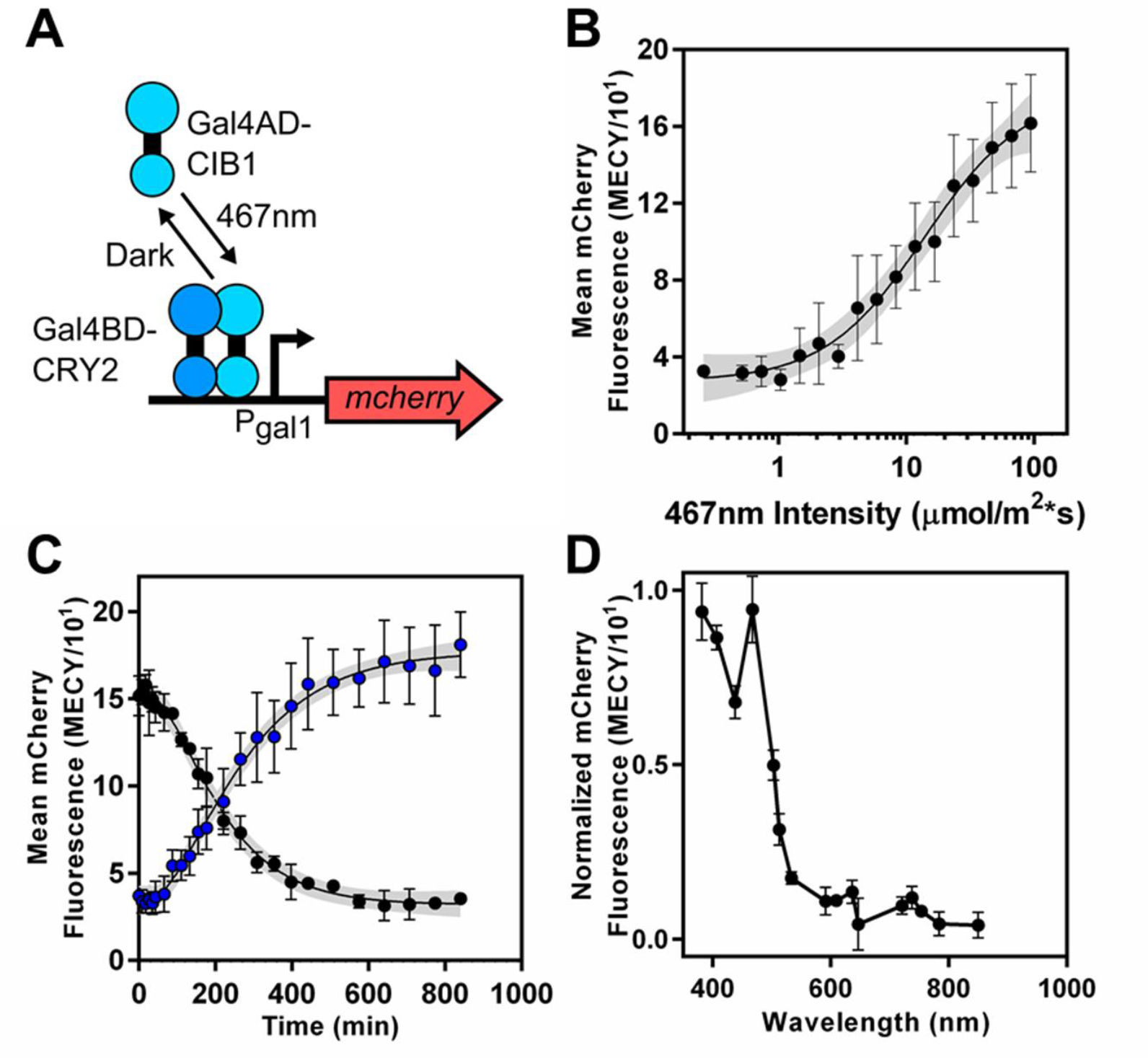
Validating the LPA with yeast by characterizing S. *cerevisiae* CRY2-CIB1 Y2H. (**a**) CRY2-CIB1 Y2H system. (**b**) 467 nm intensity transfer function. The black line represents the best fit to a Hill function. (**c**) Step activation and de-activation kinetics. Cells were either preconditioned for 4 h in the dark and switched to 88.8 μmol m^−2^ s^−1^ 467 nm (blue dots) or preconditioned in 88.8 μmol m^−2^ s^−1^ 467 nm for 10 h and switched to dark (black dots) at time zero. Black lines represent best fits (**Supplementary Tables 9** and **11**) to a kinetic model (**Methods**). (d) FAS. Experiments were performed as in **Figure 3a**, but with 88.8 μmol m^−2^ s^−1^ photon flux. Grey envelopes represent 95% confidence interval. Error bars represent the SEM of mCherry levels from three experiments over three days (**b-c**) or a single day (**d**).

To demonstrate compatibility of the LPA with yeast, we characterized the CRY2-CIB1 Y2H I/O. We outfitted the top and bottom positions of an LPA with 467 nm (blue) LEDs, and programmed them to emit intensities between 0.26 and 1057.00 μmol m^−2^ s^−1^. We grew yeast expressing CRY2-CIB1 Y2H with an mCherry output in a shaking LPA under these conditions for 18 or 24 h and measured output using flow cytometry (**Methods**). Between 0.26 and 93.50 μmol m^−2^ s^−1^, mCherry increases in a Hill-like manner with 5.6-fold dynamic range, *n_H_* of 1.3 and *k_1/2_* of 10.74 μmol m^−2^ s^−1^ (**Fig. 5b**), consistent with previous reports^13,26,54^. Though mCherry increases slightly at higher intensities, we observe growth defects, likely due to heating or phototoxicity (**Supplementary Fig. 22**).

To characterize CRY2-CIB1 Y2H dynamics, we performed a step increase and decrease in blue light and measured the mCherry response over time (**Methods**). We observe a delay followed by an exponential increase (step-on) or decrease (step-off) in mCherry to a final steady state (**Fig. 5c**). By fitting the data to a first order ODE model with a delay (**Methods**), we quantify the delay to be 75.13 min, and the time to reach 50% of the final mCherry level to be 222 min (**Supplementary Table 11**). The slower dynamics compared to CcaS-CcaR are likely due to slower rates of transcriptional activation and cell division, as well as the additional steps of mRNA processing and nuclear/cytoplasmic transport.

Next, we measured the CRY2/CIB1 Y2H FAS by outfitting the bottom position of an LPA with LEDs between 382 and 849 nm and programming their outputs to 88.80 μmol m^−2^ s^−1^, the highest intensity possible without reducing the output of the dimmest (533 nm) LED. This experiment reveals peak activation in response to the 382 nm and 467 nm LEDs and broad wavelength responsivity in the blue (>50% response from 382 to 503 nm) which matches the broad, multipeaked in-vitro absorbance spectrum of CRY2^55^ (**Fig. 5d**). We conclude that the LPA is compatible with yeast optogenetic tools.

### Spatial control of mammalian cell gene expression with PHYB/VNP-PIF6

We recently modified the surface of viral nanoparticles (VNPs) with Phytochrome Interacting Factor 6 (VNP-PIF6) to bind a heterologously expressed nuclear localization sequence tagged Phytochrome B (PHYB-NLS) and deliver a transgene to the mammalian cell nucleus in a red light activated, far-red deactivated manner^56^ (**Fig. 6a**). Using an early LPA prototype, we tuned delivery of *gfp* to HeLa cells from 35 to 600% that of wild-type virus. Using custom photomasks, we also patterned GFP expression within different regions of a confluent culture well. Delivery of red light alone resulted in low contrast between cells inside and outside the pattern. Co-delivering far red reduced expression in unwanted areas and increased contrast. However, this prototype had numerous shortcomings including a requirement for a glass bottom plate that often fractured during experiments.

**Figure 6.**
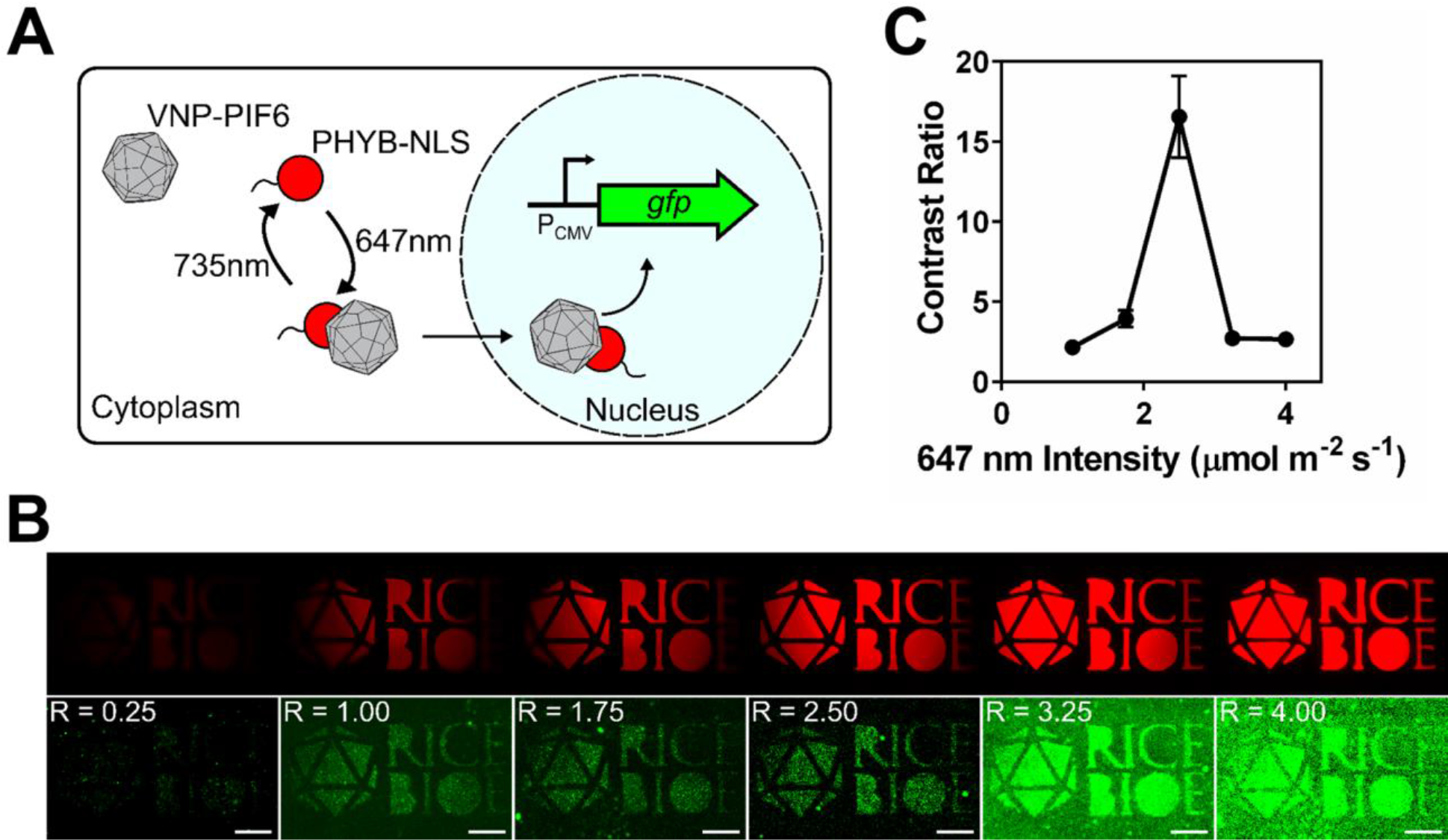
Validating the LPA with mammalian cells by spatial patterning of transgene delivery with PHYB/VNP-PIF6. (**a**) PHYB/VNP-PIF6 system. (**b**) HeLa cells expressing PhyB(908)-NLS were treated with 15μM PCB and VNP-PIF6 (2,000 viruses per cell) and exposed through a photomask (top row and **Supplementary Fig. 17b**) to 2.0 μmol m^−2^ s^−1^ far-red for 30 min followed by 1.0 μmol m^−2^ s^−1^ far-red and variable red intensities (shown in μmol m^−2^ s^−1^ in upper left of cell fluorescence images) for 60 min. Image corresponding to red intensity of 1.75 μmol m^−2^ s^−1^ was acquired with photomultiplier setting of 80 while all others were acquired with photomultiplier setting of 60. (**c**) Contrast ratio of cell fluorescence patterns from images in panel b (bottom row). Symbols represent the average ratio of pixel intensity in three light-exposed regions to an unexposed region, while error bars represent the SEM.

To validate compatibility with mammalian cells and extend our previous results, we outfitted the top and bottom positions of an LPA with 647 (red) and 735 nm (far red) LEDs and replaced the plate adapter gasket (**Supplementary Fig. 17a**) with a laser-cut black nitrile photomask (**Supplementary Fig. 17b**). We preconditioned HeLa cells in 6 wells of 1 plate with 2.00 μmol m^−2^ s^−1^ far red light for 30 min and then reduced it to 1.00 μmol m^−2^ s^−1^ while introducing variable red light between 0.25 and 4 μmol m^−2^ s^−1^ for 1 h. We allowed GFP to accumulate during a 48 h dark incubation and then imaged the resulting expression patterns by fluorescence microscopy (**Methods**). As expected, GFP expression increases with red intensity (**Fig. 6b**). Additionally, the contrast ratio (GFP intensity in exposed divided by that in non-exposed regions) reaches an optimal value of 16.6, near the maximum dynamic range of the system, at an intermediate red:far red ratio of 2.5 (**Fig. 6c**).

### Using the LPA to entrain the cyanobacterial circadian clock

The cyanobacterium *Synechococcus elongatus* PCC7942 is a model for studying circadian rhythm^57^. Exposing cells to 12 h dark/12 h light cycles entrains the circadian clock. After entrainment, a roughly 24 h transcriptional oscillation^58,59^ is maintained for roughly 30–64% of the genome, even under constant light^60^.

The time-intensive nature of the entrainment protocol makes it difficult to conduct multiple parallel experiments in order to study clock properties and one group has recently sought to simplify the procedure by using a computer-controlled LED array^51^. To examine whether the LPA can be used to automate circadian entrainment, we outfitted the top and bottom positions of three devices with 647 and 467 nm LEDs to provide a balanced photosynthetic active radiation spectrum typical of grow lights. We programmed 6 different signals across the devices wherein cells were exposed to 3 cycles starting at different times. Luminescence from a commonly used reporter system^60,61^ wherein luciferase substrate is produced from the “dusk-peaking” promoter *psbAI* and luciferase from the “dawn-peaking” promoter *kaiBC* was then measured (**Fig. 7a**). As expected, the oscillations of cultures that were entrained starting at different times showed different phases (**Fig. 7b**). The order of peaks in luminescence was also as expected, with the cultures entrained starting at 0 h (red) peaking before cultures entrained starting at 4 h (orange), 8 h (yellow), 12 h (green), 16 h (blue) and 20 h (purple) (**Fig. 7b,Fig. 7c**). However, the position of the peaks exhibits some dispersion, potentially due to heating of the LPA or the transition of cultures from the LPA to the plate reader.

**Figure 7.**
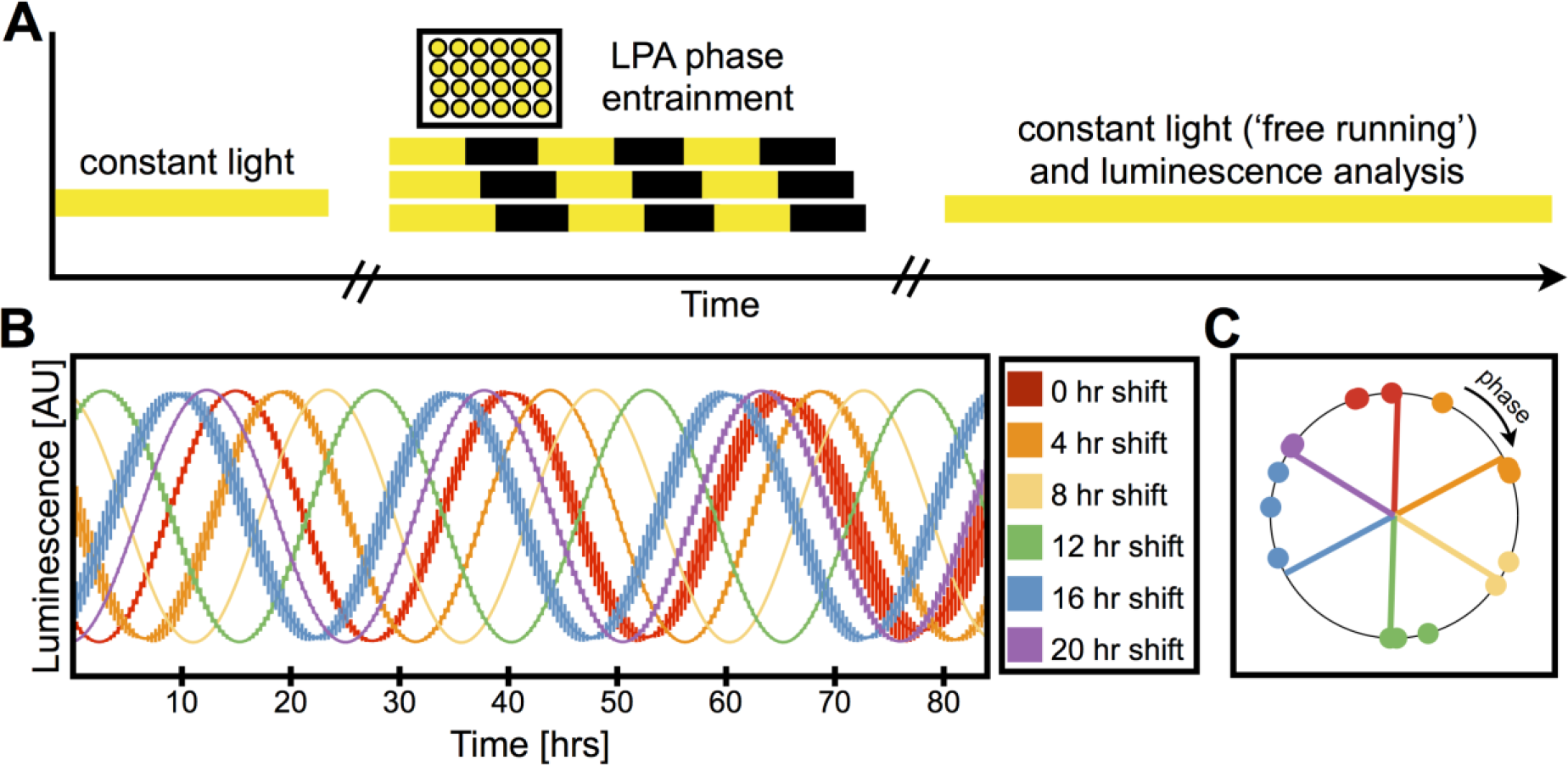
Validating the LPA with cyanobacteria by entraining circadian rhythm in S. *elongatus*. (**a**) Schematic representation of entrainment protocol. Cells grown under constant light were placed in the LPA and entrained with three light-dark cycles, with dark periods starting at different times in order to shift the phases of each well. The cells were then transferred to a plate reader for measurement of luminescence under constant light. (**b**) Luminescence oscillation for cultures entrained starting at different times. Cultures were entrained starting at zero (red), four (orange), eight (yellow), 12 (green), 16 (blue) and 20 (purple) hours. Fits of raw data were normalized to equalize peak height, and the normalized fits were averaged across three replicates except for the cultures in yellow for which only two replicates could be obtained. Bars indicate SEM. (**c**) Phase of cultures entrained starting at different times. Colors are the same as in **b**. Dots indicate cosine acrophase (phase of peak) of the replicates. Lines indicate the best fit for the expected locations of acrophase for perfect entrainment.

## Discussion

The LPA establishes new standards of flexibility, user-friendliness, and affordability in non-neural optogenetics and photobiology hardware – features that should facilitate widespread adoption. In terms of flexibility, the LPA is compatible with bacteria, yeast, and mammalian cells and other common model organisms. The ~1 mL well volume and the shaker adapters make the device compatible with most model organism growth protocols. The diversity of available LEDs theoretically makes the LPA compatible with all known genetically encoded photoreceptors and most photobiological processes. The second LED position allows improved spatial and dynamical control of photoreversible optogenetic tools and characterization of reverse action spectra. Different spatial patterns can easily be applied by using laser cut gaskets. In addition to these features, the LPA retains the high standards of programmability and experimental precision of the LTA^47^.

The ease of assembly, intuitive Iris interface, simplified liquid handling, and automated calibration script make the LPA user friendly. The LPA can be assembled easily and quickly by a non-expert using fully documented assembly instructions and tutorial videos. Iris converts high-level light function specifications into low level hardware operations, greatly reducing programming requirements. LPA samples can be loaded and harvested using a multichannel pipettor, and the plate can be sealed and unsealed with an adhesive lid in a single step, facilitating high throughput experimentation. The plate is also compatible with plate readers and plate-based flow cytometers, enabling direct measurements of absorbance, fluorescence, or luminescence without additional liquid handling. Finally, our MATLAB script simplifies calibration for experiments where the same LED is used in multiple wells.

The LPA is more economical than comparable devices. The 24-well plate costs $7.70 and can be reused dozens of times, lowering per sample cost to <$0.01. Furthermore, the $400 component cost drops to $150 with access to a 3D printer. These low costs make the LPA practical for large scale experiments (**Supplementary Fig. 23**).

The action spectra of optogenetic tools are seldom measured due to the lack of convenient hardware. Using the LPA, both the FAS and RAS can be measured with 24 wavelength resolution in one experiment. This feature should permit improved characterization of light responsive signaling pathways and enable cross-talk to be minimized when using multiple optogenetic photoreceptors.

Despite early efforts^62,63^, there is little data directly comparing the performance features of optogenetic tools. Accordingly, researchers may have difficulty selecting the best tool for a given experiment. On the other hand, the LPA can be used to directly compare optogenetic tools, including those in different organisms. For example, in this study we found that CRY2/CIB1 Y2H is 50 times less sensitive to light than CcaS-CcaR (compare *k_1/2_* in **Figs. 3b, Figs. 5b**). Using the basic protocols established here, the LPA could be used to quantitatively compare the spectral, intensity dependent, and dynamical properties of all published optogenetic tools with gene expression outputs in bacteria, yeast, mammalian cells and other organisms, providing much needed information to the community that should accelerate the adoption of optogenetics.

Many circadian assays, such as those measuring the sensitivity of the rhythm to optical perturbations as a function of time of day, are onerous and present a significant barrier to the researcher. The LPA provides a compact and economical solution to this problem. Additionally, the ability to program two wavelengths may ease studies of organisms and processes that respond differently to different wavelengths such as phase resetting in the green algae *C. reinhardtii* in which shifts in the circadian rhythm are wavelength dependent as a function of cellular status. The ability to program 24 light signals simultaneously may also ease experiments such as testing growth of mutants with longer or shorter circadian periods with different lengths of light and dark^6466^.

By releasing the LPA and Iris with an open-source license, we aim to incorporate other researchers as developers to extend and improve the platform. Similar to a recent example ^67^ we have made all design files, a bill of materials, and protocols needed to fabricate, assemble, and operate the device, as well as standard format source code on a public version control system freely available. From this repository, the community can download up-to-date file versions, suggest or make improvements or modifications, design new accessories, and upload all updates for others to use and build upon further. Future designs could incorporate more LEDs per well, sample injection, and on-board sensors for measurement of absorbance, fluorescence, or luminescence, enabling optogenetic feedback control^68^. Better heat dissipation for high intensity experiments could be achieved through addition of heat sinks, fans, chassis parts with better ventilation, or even active temperature feedback control. 96-well or greater plate format designs should also be possible. With 3D printing and microcontroller technologies becoming increasingly inexpensive and accessible, such advances can be made by researchers without formal electrical and mechanical engineering backgrounds. We hope that this work not only enables new researchers to perform optogenetics and photobiology experiments, but also motivates them join the growing open-hardware community.

## Methods

### LPA

Detailed descriptions of LPA design, fabrication, assembly, calibration, and operation, and CAD files and firmware are included in **Supplementary Information**. The most up-to-date documentation and file versions can be found at *taborlab.rice.edu/hardware*.

### Iris

Source code and detailed descriptions of the staggered start algorithm, waveform handling, file specifications, randomization and de-randomization procedure, and LPF creation using Python are included in **Supplementary Information**. The most up-to-date documentation and source code can be accessed at *taborlab.rice.edu/software*.

### CcaS-CcaR

To make frozen starter aliquots, a 3 mL LB + 50 μg/mL kanamycin, 100 μg/mL spectinomycin and 34 μg/mL chloramphenicol culture was inoculated from a −80°C stock and grown (37°C, 250 rpm) to OD600 < 0.3. 30% glycerol was added, absorbance measured, and 50μL aliquots made and stored at −80°C. For experiments, M9 + antibiotics was inoculated with a starter aliquot, to OD_600_ = 0.00015. 500 μL starter culture was added to each well of the 24-well plate and the plate was sealed with adhesive foil.

The plate was placed into the LPA and the assembly mounted on a shaker/incubator at 37μC, 250 rpm for 8 h. The plate was then removed and chilled in an ice-water bath. Cells were chilled for ≥ 15 min and the foil removed. 200 μL from each well was transferred to flow cytometry tubes containing 1 mL PBS, and cells were treated with rifampicin as before^47^.

### CRY2-CIB1 Y2H

A 3 mL SD (-Ura, -Trp, -Leu) medium starter culture was inoculated from a −80°C stock and grown (30°C, 250 r.p.m) overnight to a final density of approximately OD_600_ = 2. The starter culture was diluted into fresh SD to OD600 = 0.001. The plate was prepared and loaded/unloaded as with *E. coli*. Cells were grown in the LPA (30°C, 250 rpm) for 24 h (step-off dynamics experiments) or 18 h (other experiments). Samples were harvested as described for *E. coli* without rifampicin treatment.

### Flow cytometry

Cytometry was performed with a BD FACScan flow cytometer as previously^47^. Settings used for each experiment are listed in **Supplementary Table 12**. Typical count rates of 1,000-2,000 events/s were used and approximately 20,000 events were captured for each sample. Calibration beads (RCP-30-5A, Spherotech) were measured at each session. Data was processed and sfGFP and mCherry fluorescence were calibrated to Molecules of Equivalent Fluorescein (MEFL) and Molecules of Equivalent Cy5 (MECY) with FlowCal^69^.

### CcaS-CcaR Model

Data was modeled as previously^47^. Parameter fitting was performed in GraphPad. All parameters were constrained to > 0, and *τ_delay_* and *k_g_* are shared between step-on and step-off data sets.

### CRY2-CIB1 Y2H Model

We apply a first order linear ODE model

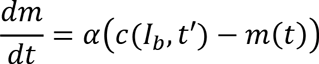

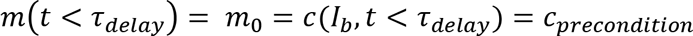

where the rate of change of mCherry, *m*(*t*), is a function of the difference between the mCherry “set-point”, *c*(*t′*) and the current mCherry concentration. *c*(*t′*) is blue light intensity, *I_b_*, mapped to mCherry units through the steady-state transfer function at when the light signal last changed, *t′* = *t* − *τ_deiay_*. At *t* < *τ_delay_*, mCherry is equal to its initial value, which is determined by the preconditioning set-point, *c_precondition_*.

The solution to this ODE has the following piece-wise form:

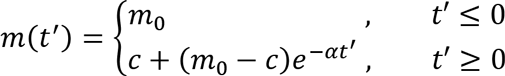

where mCherry remains at its initial value, *m_0_*, for *t* ≤ *τ_delay_*, and exponentially increases or decreases with rate constant *α* for *t* ≥ *τ_delay_*.

Parameter fitting was performed by nonlinear least squares in GraphPad. All parameters were constrained to > 0, and *τ_delay_* and *α* were shared between step-on and step-off data sets.

### PH YB/VNP-PIF6

Experimental protocols were as described^56^ except that three sheets of diffuser paper (#3008, Rosco) were used in place of the LED spacer gasket. Adhesive foil was not used to avoid interference from reflected light.

### Synechococcus Native Circadian Rhythm

1.5ml of a culture of AMC462^70^ growing in exponential phase diluted to OD_750_ 0.05 was pipetted into each well of the 24-well cell culture plate. Extra distilled water was pipetted between wells to reduce evaporation, and the plate was covered with a transparent lid (Visiplate 24-TC Part: 1450-6045) with adhesive foil (F96VWR100). The cells were then grown at 30° C with shaking at 255 rpm for 4 days.

### Luminescence data acquisition and analysis

Luminescence of 150 μl of culture in a 96-well black with clear bottom plate with transparent lid was measured using a Tecan M200 with a luminescence module and an integration time of 1000 ms. Between readings, the plate was moved out of the machine for 20 min and kept under a fluorescent lamp at 4500 lux. Readings were taken for a total of 160 h.

Luminesce data was fitted with BRASS (Biological Rhythms Analysis Software System; A. J. Millar laboratory, University of Edinburgh, Scotland, United Kingdom) and FFT-NLLS (Fast Fourier Transform-Nonlinear Least Squares)^71,72^. At least 72 h of each culture was included in the analysis.

The best fits for the expected phases were calculated by minimizing the root mean square distance from the location of the expected phase to the location of experimental phase using the GRG Nonlinear solving method in Excel.

## Acknowledgements

We thank M. McClean for providing yMM1146 and yMM1081, S. Golden for *Synechococcus* AMC462, and M. Romer for assistance with *Synechococcus* experiments. This work was supported by the ONR (MURI N000141310074), NSF (EFRI-1137266), and NIH (1R21AI115014-01A1). B.L. is supported by an NDSEG Fellowship. L.H. was supported by NASA NSTRF NNX11AN39H. E.J.G. is supported by a Ford Foundation Fellowship. The cyanobacterial work was supported by a DOE Office of Science Early Career Research Program (DE-SC0006394) to D.S. and R.Y. is a Simons Foundation Fellow of the Life Sciences Research Foundation.

## Author Contributions

K.G., E.O., and J.T. conceived of the project. K.G. (chassis and auxillary parts), E.O. and S.C. (circuit board) designed the LPA. K.G, E.O., and P.R. built LPAs. S.C. designed and wrote the LPA firmware and calibration software. L.H., B.L., and F.E. designed and wrote Iris with equal contributions. K.G., E.O., and J.T. (*E. coli, S. cerevisiae* and mammalian) and R.Y. and D.S. (cyanobacteria) designed experiments. K.G. (*E. coli, S. cerevisiae*), E.G. (mammalian), and R.Y. (cyanobacteria) performed experiments, and K.G. and R.Y. analyzed results. J.T., K.G., E.O., L.H., S.C., R.Y. and D.S. wrote the manuscript. J.S., D.S. and J.T. supervised the project.

## Material and Correspondence

Correspondence and material requests should be addressed to J.J.T.

## Supplementary Information

Documentation and procedures for the LPA and Iris, best-fit parameters, cytometry settings, phototoxicity and/or LPA heating measurements are included in Supplementary Information. CAD files, firmware, Iris source code, image analysis calibration script, example calibration files, and supplementary videos are included in Supplementary Files.

